# Ferric Citrate Uptake is a Virulence Factor in Uropathogenic Escherichia coli

**DOI:** 10.1101/2021.11.12.468464

**Authors:** Arwen E. Frick-Cheng, Anna Sintsova, Sara N. Smith, Ali Pirani, Evan S. Snitkin, Harry L.T Mobley

## Abstract

More than half of women will experience a urinary tract infection (UTI) with uropathogenic *Escherichia coli* (UPEC) causing ∼80% of uncomplicated cases. Iron acquisition systems are essential for uropathogenesis, and UPEC encode functionally redundant iron acquisition systems, underlining their importance. However, a recent UPEC clinical isolate, HM7 lacks this functional redundancy and instead encodes a sole siderophore, enterobactin. To determine if *E. coli* HM7 possesses unidentified iron acquisition systems, we performed RNA-sequencing under iron-limiting conditions and demonstrated that the ferric citrate uptake system (*fecABCDE* and *fecIR*) was highly upregulated. Importantly, there are high levels of citrate within urine, some of which is bound to iron, and the *fec* system is highly enriched in UPEC isolates compared to environmental or fecal strains. Therefore, we hypothesized that HM7 and other similar strains use the *fec* system to acquire iron in the host. Deletion of both enterobactin biosynthesis and ferric citrate uptake (Δ*entB*/Δ*fecA*) abrogates use of ferric citrate as an iron source and *fecA* provides an advantage in human urine in absence of enterobactin. However, in a UTI mouse model, *fecA* is a fitness factor independent of enterobactin production, likely due to the action of host Lipocalin-2 chelating ferrienterobactin. These findings indicate that ferric citrate uptake is used as an iron source when siderophore efficacy is limited, such as in the host during UTI. Defining these novel compensatory mechanisms and understanding the nutritional hierarchy of preferred iron sources within the urinary tract are important in the search for new approaches to combat UTI.

**Importance:** UPEC, the primary causative agent of uncomplicated UTI is responsible for five billion dollars in healthcare costs in the US each year. Rates of antibiotic resistance are on the rise therefore it is vital to understand the mechanisms of UPEC pathogenesis to uncover potential targets for novel therapeutics. Iron acquisition systems used to obtain iron from sequestered host sources are essential for UPEC survival during UTI and have been used as vaccine targets to prevent infection. This study reveals the ferric citrate uptake system is another important iron acquisition system that is highly enriched in UPEC strains. Ferric citrate uptake has not previously been associated with these pathogenic isolates, underlining the importance of the continued study of these strains to fully understand their mechanisms of pathogenesis.

## Introduction

More than half of women will experience a urinary tract infection (UTI) during their lifetime, and 25% of infections recur (1, 2) with uropathogenic *Escherichia coli* (UPEC) causing 80% of uncomplicated cases (3, 4). These infections are responsible for an annual five billion dollars of health care costs in the U.S. alone (5, 6) To survive within the host UPEC encodes a wide array of virulence factors that include toxins, adhesins and iron acquisition systems (6-9). Iron is an essential cofactor for many biological processes including DNA replication, DNA repair, and central metabolism (10, 11). Consequently, mammalian hosts employ “nutritional immunity” wherein iron is sequestered within proteins or molecules such as transferrin, lactoferrin, ferritin, and hemoglobin and is not readily accessible to bacteria (12, 13). To survive in the host, UPEC has evolved mechanisms to acquire iron from these sequestered sources which fall into two broad categories: heme receptors and siderophores. Heme receptors import heme, allowing the bacteria to utilize the bound iron while siderophores are small molecules with extraordinarily high affinities for iron (K_d_ ranging from 10^23^ to 10^52^ M^-1^) (14, 15), which allow them to strip iron from sequestered sources.

UPEC can encode up to five iron acquisition systems: heme receptors (ChuA and Hma), and four siderophores (enterobactin, salmochelin aerobactin, and yersiniabactin) (16-18). UPEC strains often employ a subset of these systems. For example, prototypical UPEC strain CFT073 encodes heme receptors and produces enterobactin, salmochelin and aerobactin. This high level of functional redundancy is essential for UPEC survival within the host due in part to specific host defenses. For example, the innate immune protein Lipocalin-2 (Lcn2) binds ferric and aferric enterobactin, preventing the bacterium from utilizing this siderophore (19). Therefore, UPEC cannot rely upon a single method of iron acquisition.

While heme receptors and siderophores are traditional iron acquisition systems utilized by UPEC and other pathogenic bacteria, there are other methods. For instance, citrate is a weak iron chelator and ferric citrate complexes can be imported through the ferric citrate transporter, or *fec* system (20, 21). A study investigating *E. coli* strains that caused bovine mastitis (MPEC) discovered that *fec* was a major pathogenic determinate of these strains; in 62 MPEC strains, ∼98% encoded the *fec* system (22). The high citrate levels in milk (∼10 mM) provide a pool of ferric citrate for these bacterial strains to use as an iron source via the *fec* system, allowing MPEC to grow in milk and induce mastitis (22).

Overall, little has been done to define the role of the *fec* system in the context of pathogenesis. However, there is a substantial body of work defining its regulation and mechanism of action. The *fec* system is composed of two operons, *fecIR* and *fecABCDE* (23-25). *fecIR* is Fur-regulated and expressed under iron-limiting conditions (26), while *fecABCDE*, is specifically transcribed via FecI, an alternative sigma factor, when ferric citrate is present (21, 26). FecA is a TonB-dependent outer membrane receptor, and FecBCDE comprise an ABC transporter (23, 24, 27).

In this study we have investigated the redundancy of iron acquisition systems in a collection of UPEC strains that caused symptomatic UTI in healthy, college-aged women (28, 29). One of these clinical isolates, HM7, lacked the functional redundancy in iron acquisition systems characteristic of most UPEC strains and only encoded a sole siderophore, enterobactin. While this strain lacks the traditional methods of iron acquisition and has no clear mechanism to prevent Lcn2 from inactivating enterobactin, it is clearly pathogenic as it was isolated from a young woman with cystitis and was present in the urine at ≥10^5^CFU/mL. In this study, we sought to determine how this novel strain acquires iron from the host. First, we empirically demonstrated that Lcn2-susceptible enterobactin is the sole siderophore produced by HM7. Then, using RNA-sequencing (RNA-seq), we found the ferric citrate uptake system highly upregulated under iron limitation. Furthermore, we discovered that the *fec* system is highly enriched in UPEC isolates when compared to fecal or environmental strains, and that there is a small, but significant cohort of UPEC strains that encode a single siderophore. Additionally, HM7 can use ferric citrate as an iron source through the *fec* system and enterobactin *in vitro*, and the *fec* system is a fitness factor *in vivo*. Our study characterizes ferric citrate uptake as a UPEC virulence factor, adding a novel iron-scavenging mechanism that UPEC uses to survive within the urinary tract.

## Results

### Clinical UPEC isolate HM7 lacks all but one of the iron acquisition systems associated with UPEC

Roughly, there are up to five systems that UPEC use to acquire iron from the host (**Fig. 1A**). Most UPEC strains encode four of the five systems, including the three major UPEC type strains (CFT073, UTI89, and 536, **Fig. 1A, B**). However recent clinical isolate HM7 (28) encodes a single system, enterobactin, (**Fig. 1A, B**). After analyzing 487 publicly available UPEC strains on the bioinformatics resource PATRIC (30) (**Table S1**), 44 strains shared the same profile as HM7, indicating that HM7 is not an outlier, and potentially represents a previously unrecognized subset of UPEC strains (**Fig. 1B**).

**Figure 1:**
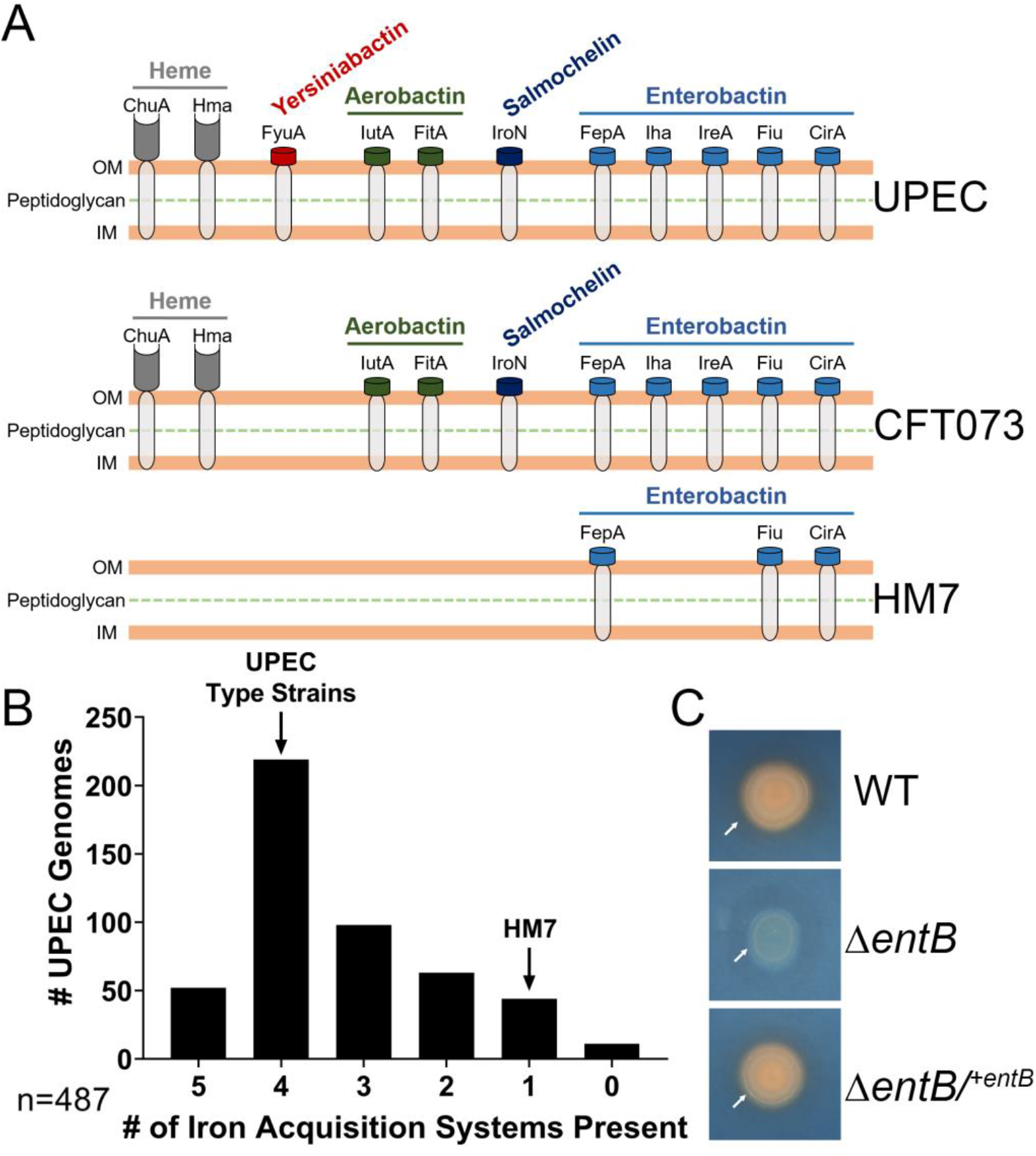
Clinical UPEC isolate HM7 encodes a single iron acquisition system. (A) Models of siderophores, siderophore uptake receptors and heme receptors in UPEC. “UPEC” indicates all known systems that have been found in UPEC while “CFT073” and “HM7” illustrate the systems in each of these indicated strains. (B) The number of iron acquisition systems present in cohort of 487 UPEC strains on the bioinformatics resource PATRIC. The five systems are composed of heme uptake (ChuA or Hma), and four siderophores (enterobactin, salmochelin, aerobactin and yersiniabactin). Presence was determined by at ≥80% protein identity and coverage of select genes for each system: heme uptake (*chuA* or *hma*), enterobactin (*entB*), salmochelin (*iroB*), aerobactin (*iucA*), and yersiniabactin (*irp1*). Genes selected for siderophores are all involved in biosynthesis. 11% of strains have five systems, 45% of strains have four, 20% of strains have three, 13% of strains have two, 9% of strains have one, and 2% appear to have none. (C) Siderophore production assayed through growth on CAS agar. 5 uL of an overnight LB culture were spotted on CAS agar and grown overnight at 37°C. A change from blue to orange indicates siderophore activity. White arrow indicates the colony in all three strains, and the orange halo in WT and complemented strain is due to diffusion of secreted siderophore.

### HM7 encodes a single siderophore

It was not clear how HM7 acquired iron and survived in the host since the innate immune protein Lcn2 renders enterobactin unusable by bacteria (19). Therefore, we hypothesized that HM7 encodes a novel siderophore to acquire iron. To test this, we deleted the gene *entB* (Δ*entB*), which is sufficient to disrupt enterobactin production (17), and tested siderophore production by culturing the deletion mutant on Chrome Azurol S (CAS) agar, a colorimetric iron chelation assay (31). A color change from blue to orange on the plate indicates iron chelation, which is observed on the colony itself as well as a halo around the colony due to diffusion of secreted siderophores. The wildtype (WT) strain showed robust siderophore activity that was absent in the Δ*entB* mutant but was subsequently restored by genetic complementation (Δ*entB/*^*+entB*^) (**Fig. 1C**). These results indicate that enterobactin is the sole siderophore system in HM7.

### Ferric citrate uptake is significantly upregulated during iron restriction

HM7 did not make a novel siderophore, therefore, we predicted that it might utilize a previously undiscovered or understudied iron acquisition system. To identify a list of candidate genes, we used RNAseq to determine the iron regulon of HM7. We added increasing amounts of the iron-specific chelator 2,2’-dipyridyl (Dip) to minimal M9 medium supplemented with 0.4% glucose to define an iron-restricted condition. The addition of 150 μM Dip to the base medium was sufficient to modestly limit growth due to iron restriction without introducing a severe growth defect (**Fig. S1A**). Based on previous literature (32, 33) M9 supplemented with 36 μM FeCl_3_ was the iron-replete condition (**Fig. S1A**). We confirmed these conditions reflected iron-restricted and iron-replete conditions through qRT-PCR; the iron-regulated gene *entF* was significantly and highly upregulated in the iron-restricted condition when compared to iron-replete (**Fig. S1B**).

HM7 was cultured to mid-log phase under these conditions in biological triplicate, its RNA isolated and sequenced. 368 genes were significantly downregulated in the iron-depleted condition (**Table S2**), while 393 genes were significantly upregulated (**Table 1, Table S2**). As expected, we observed that the genes for enterobactin biosynthesis and uptake as well as genes associated with iron-starvation (*nrdEFH* (34)) were upregulated (**Table 1**). Two transport systems related to iron were significantly upregulated. One was *mntH* which takes up both Mn^2+^, and Fe^2+^, although with a preference for Mn^2+^ (35). The other system was ferric citrate uptake, which is composed of two operons, *fecIR*, encoding the system’s regulatory element and σ factor, and *fecABCDE* encoding the outer membrane receptor and transport elements (**Fig. 2A**). Interestingly, *fecD* was not significantly upregulated. Unlike *mntH*, the *fec* system takes up Fe^3+^ which is dominant form of iron in the urinary tract as opposed to Fe^2+^. Furthermore, citrate is present is extremely high levels in the urinary tract, normal levels in healthy individuals vary from 1.7-6.6 mM (36). Given that MPEC uses ferric citrate in bovine milk, and the citrate concentration in milk (∼10mM) is comparable to the concentration in urine, we hypothesized that UPEC is using a similar mechanism in the urinary tract.

**Table 1:**
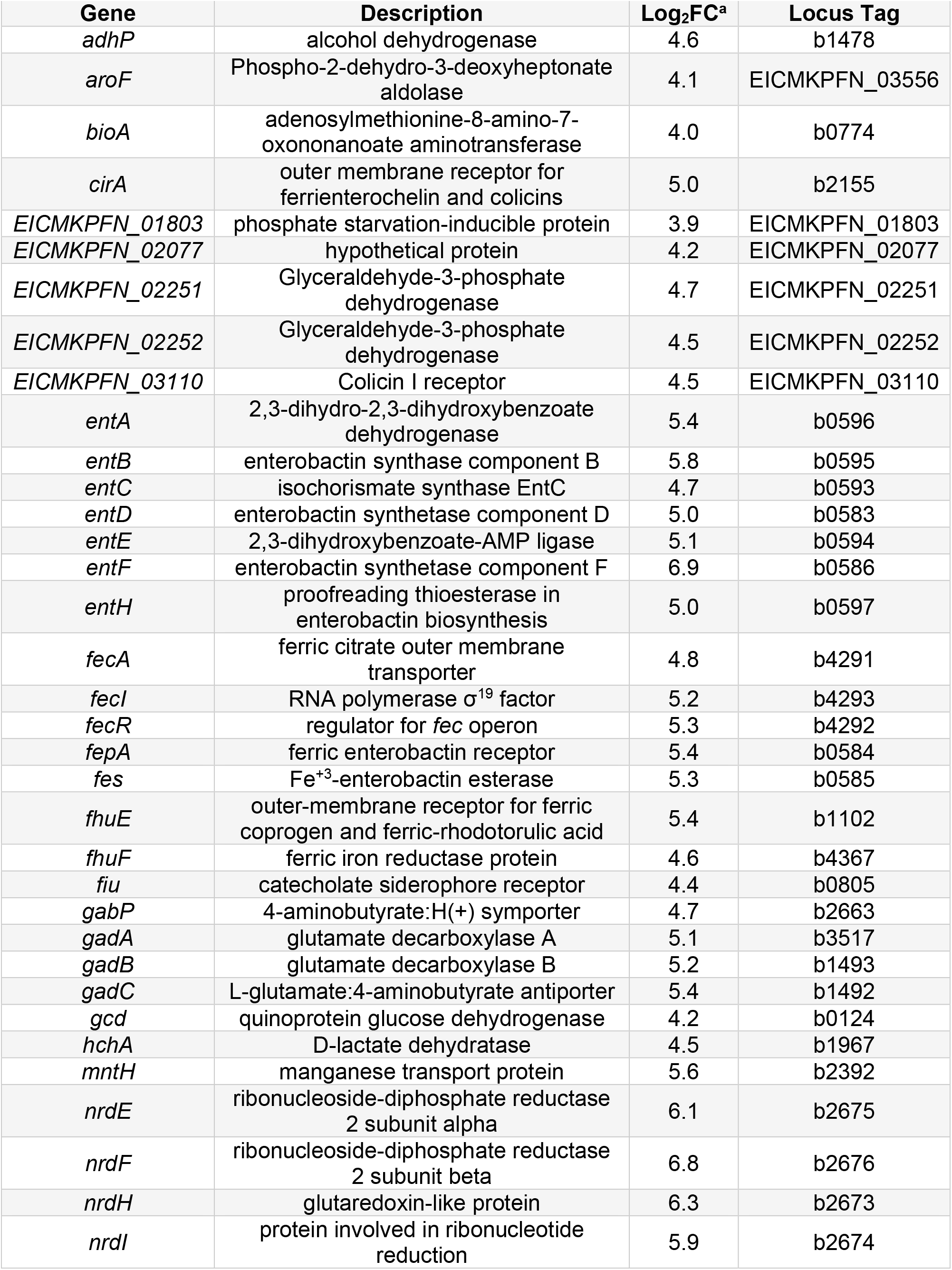
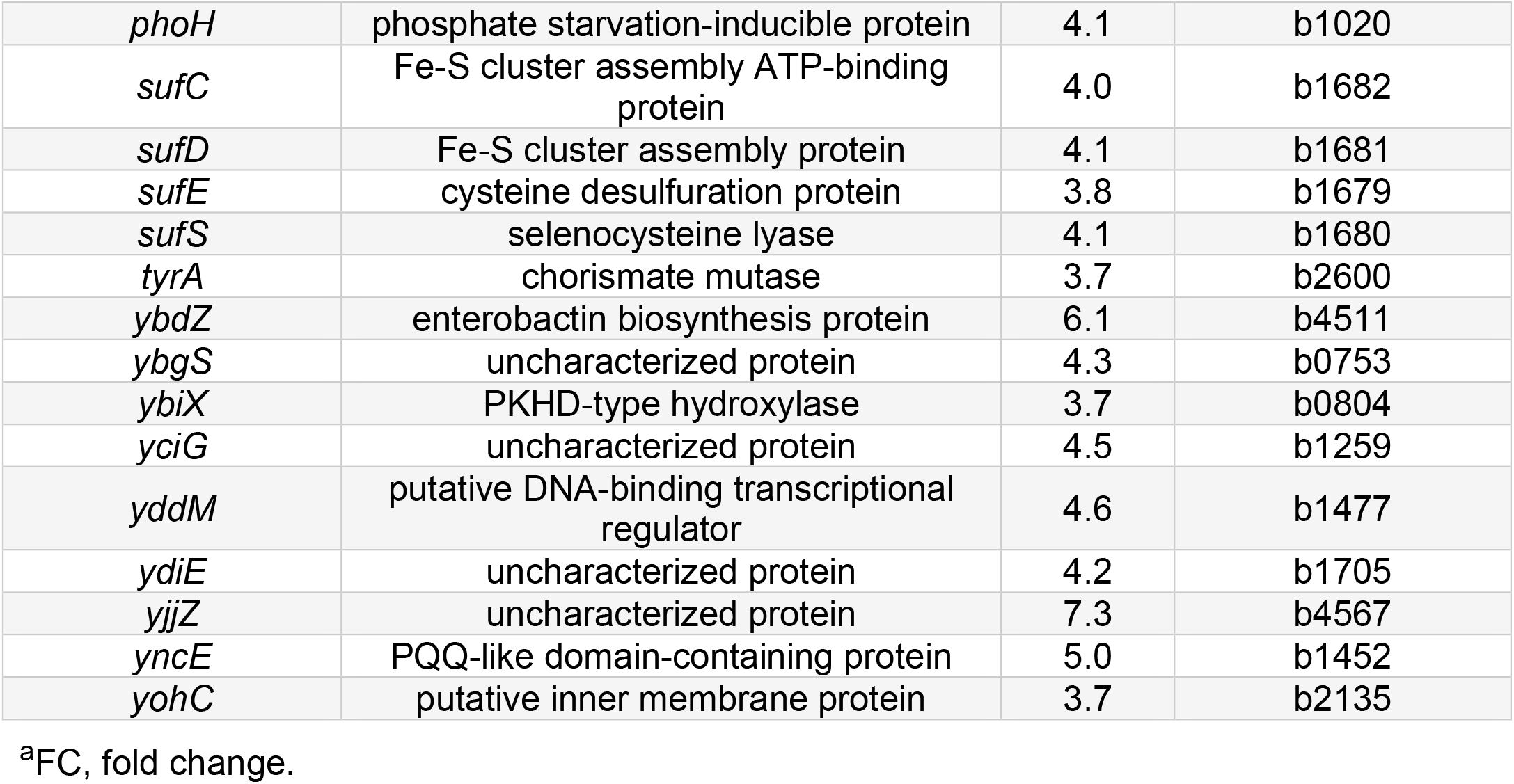
Top 50 Significantly Upregulated Genes Under Iron Limitation

**Table 2:** List of Strains and Plasmids.

| Strain | Genotype/Description | Reference/Source |
| --- | --- | --- |
| HM7 | Wild-type cystitis causing uropathogenic *E. coli* strain, isolated from a healthy young woman in 2012 | (28) |
| $\Delta entB$ | HM7 *entB*::kan, Kan<sup>r</sup> | This study |
| $\Delta fecA$ | HM7 *fecA*::cam, Cam<sup>r</sup> | This study |
| $\Delta fecA/\Delta entB$ | HM7 *fecA*::cam/*entB*::kan, Cam<sup>r</sup>, Kan<sup>r</sup> | This study |
| Plasmid | Description | Reference/Source |
| pGEN eV | Low copy number, promoterless plasmid, Spec<sup>r</sup> | This study |
| pBAD eV | pBAD-*Myc*/His A, low copy number, arabinose inducible plasmid, Amp<sup>r</sup> | Thermo-Fisher |
| pGEN *entB* | *entB* with native promoter, cloned from HM7 via Gibson assembly, Spec<sup>r</sup> | This study |
| pBAD *fecABCDE* | *fec* operon (*fecABCDE*) cloned from HM7 inserted into MCS via Gibson assembly, Amp<sup>r</sup> | This study |
| pGEX-4T-3 LCN | Human Lipocalin-2 glutathione S-transferase (GST) fusion protein, Amp<sup>r</sup> | (53) |

**Table 3:**
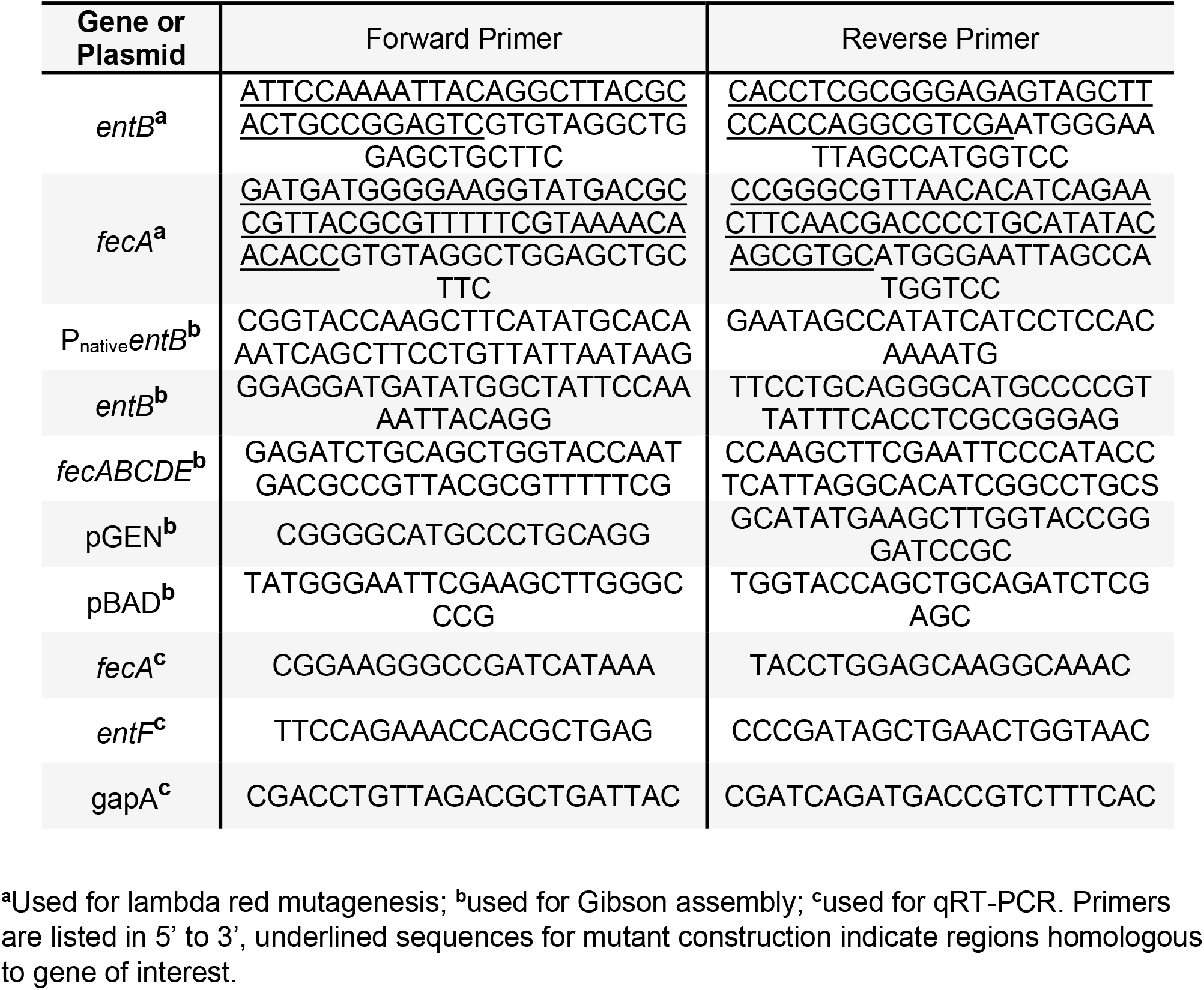
Primers used in study.

**Figure 2:**
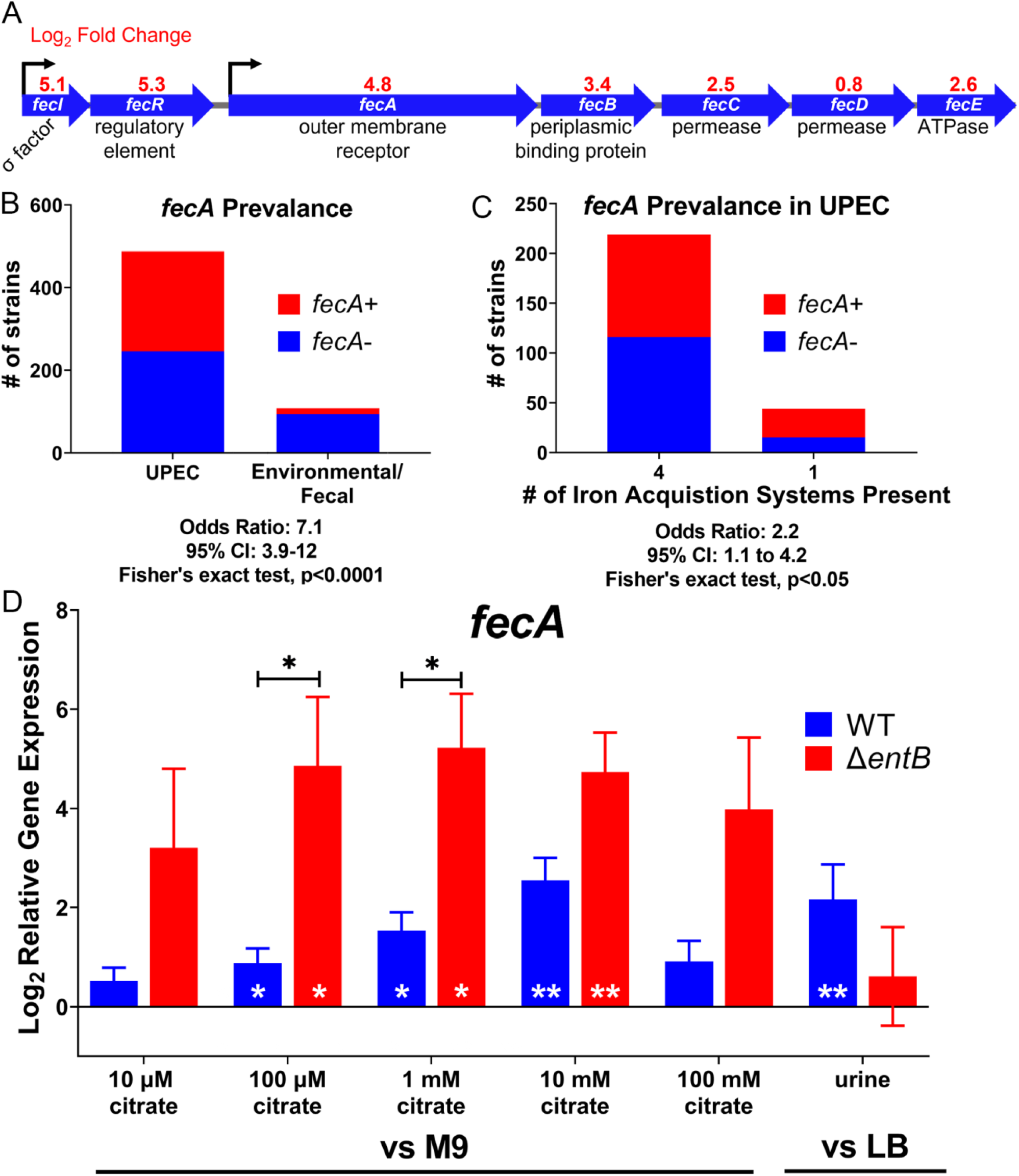
Ferric citrate uptake is a potential iron acquisition system in UPEC. (A) RNAseq revealed the ferric citrate uptake system (*fecABCDE* and *fecIR*) is upregulated in WT HM7 under iron limitation (M9 supplemented with 36μM FeCl_3_ *versus* M9 with 150 μM 2,2 dipyridyl). (B) *fecA* is enriched in UPEC strains compared to *E. coli* fecal or environmental isolates. 487 UPEC strains and 107 fecal or environmental strains were analyzed; presence of *fecA* was determined by ≥80% protein identity and coverage. (C) *fecA* is enriched in UPEC strains with a single traditional iron acquisition system (“HM7-like”), compared to strains with four traditional iron acquisition systems. (D) Gene expression of *fecA* in HM7 in either M9 medium with 0.4% glucose supplemented with increasing amounts of citrate, or in pooled human urine. Gene expression was assayed through qRT-PCR. Bars are the average of five (WT) and four (Δ*entB*) biological replicates, bars are mean, error bars are ±SEM. Black asterisks compare gene expression between WT and the Δ*entB* mutant using mixed-effects analysis with Sidak’s multiple test correction, * p<0.05. White asterisks indicate significant upregulation, determined by one sample t-test, * p<0.05, ** p<0.005.

### *fecA* is highly prevalent UPEC strains

We wanted to establish the prevalence of the *fec* system in UPEC strains, since three UPEC type strains, CFT073, UTI89 and 536 lack the *fec* system (**Fig. 1B**). When we interrogated the cohort of 487 UPEC strains, we found that ∼50% of them encoded the outer membrane receptor *fecA*, compared to only ∼12% in 107 fecal or environmental *E. coli* isolates (**Fig. 2B**). This is a highly significant association, with an odds ratio of 7.1, supporting the hypothesis of ferric citrate uptake as a UPEC virulence factor. Interestingly, this enrichment seems to be even more profound in UPEC strains with a single iron acquisition system; ∼65% of these “HM7-like” strains also encoded *fecA* compared to ∼47% of strains with four traditional iron acquisition systems (**Fig. 2C**).

### *fecA* is responsive to physiologically relevant levels of citrate

HM7 is a mostly uncharacterized clinical isolate, therefore we wanted to determine if the *fec* system is fully functional, and responsive to citrate at physiologically relevant levels. We cultured WT HM7 in M9 medium with glucose as a carbon source and supplemented with concentrations of citrate ranging from 10 μM up to 100 mM, which encompasses urinary citrate levels in a healthy population (36). We quantified *fecA* gene expression compared to M9 without citrate. We observed significant upregulation at 100 μM, 1 mM, and 10 mM citrate (**Fig. 2D**) and importantly, some of the strongest upregulation occurred at physiologically relevant concentrations (1 mM and 10 mM citrate). We also tested *fecA* expression in *ex vivo* urine pooled from healthy female volunteers, compared to expression in LB. *fecA* was significantly upregulated (**Fig. 2D**) in this physiologically relevant medium.

### *fecA* is more highly upregulated in the absence of enterobactin

Next, we wanted to determine if ferric citrate uptake could act as a compensatory mechanism in the absence of enterobactin, indicating the strain can use ferric citrate as an alternative iron source. Accordingly, we repeated the citrate sensitivity experiments using the Δ*entB* mutant. Significant upregulation at 100 μM, 1 mM, and 10 mM citrate was recapitulated in the mutant strain (**Fig. 2D**). Furthermore, at both 100 μM and 1 mM citrate the Δ*entB* mutant had significantly higher expression of *fecA* compared to WT, a phenomenon that was trending in all concentrations of citrate. Interestingly, while expression of *fecA* dropped at 100 mM citrate in WT, it remained highly elevated in the Δ*entB* mutant, indicating that perhaps enterobactin is the preferred mechanism for iron acquisition, but in its absence, the *fec* system can be utilized. These results support the hypothesis that HM7 is using ferric citrate as an iron source, especially in the absence of Fe^3+^ uptake by siderophores.

### HM7 uses ferric citrate as an iron source through the *fec* system or enterobactin

To determine if HM7 can use ferric citrate as an iron source, we added high levels (100 mM) of citrate to M9 mediums so most of the iron would be complexed within citrate. The bacteria have two ways to acquire iron: either enterobactin will chelate iron from ferric citrate, or the *fec* system will import ferric citrate. To nullify ferric citrate uptake, we deleted the outer membrane receptor gene, *fecA* (Δ*fecA*). We also constructed a double mutant (Δ*fecA*/Δ*entB*). With these assumptions, only the double mutant, Δ*fecA*/Δ*entB*, would have a growth defect at high citrate concentrations, since the Δ*fecA* mutant could still utilize enterobactin, and the Δ*entB* mutant could still utilize the *fec* system. As expected, only the Δ*fecA*/Δ*entB* mutant had a profound growth defect with the addition of 100 mM citrate (**Fig. 3Aii**) while none of the mutants had a growth defect in LB or M9 medium alone (**Fig. S2, Fig 3Ai**). This is an iron-specific defect since chemical complementation with 1 mM FeCl_3_ rescued the growth of the double mutant (**Fig. 3Aiii**).

**Figure 3:**
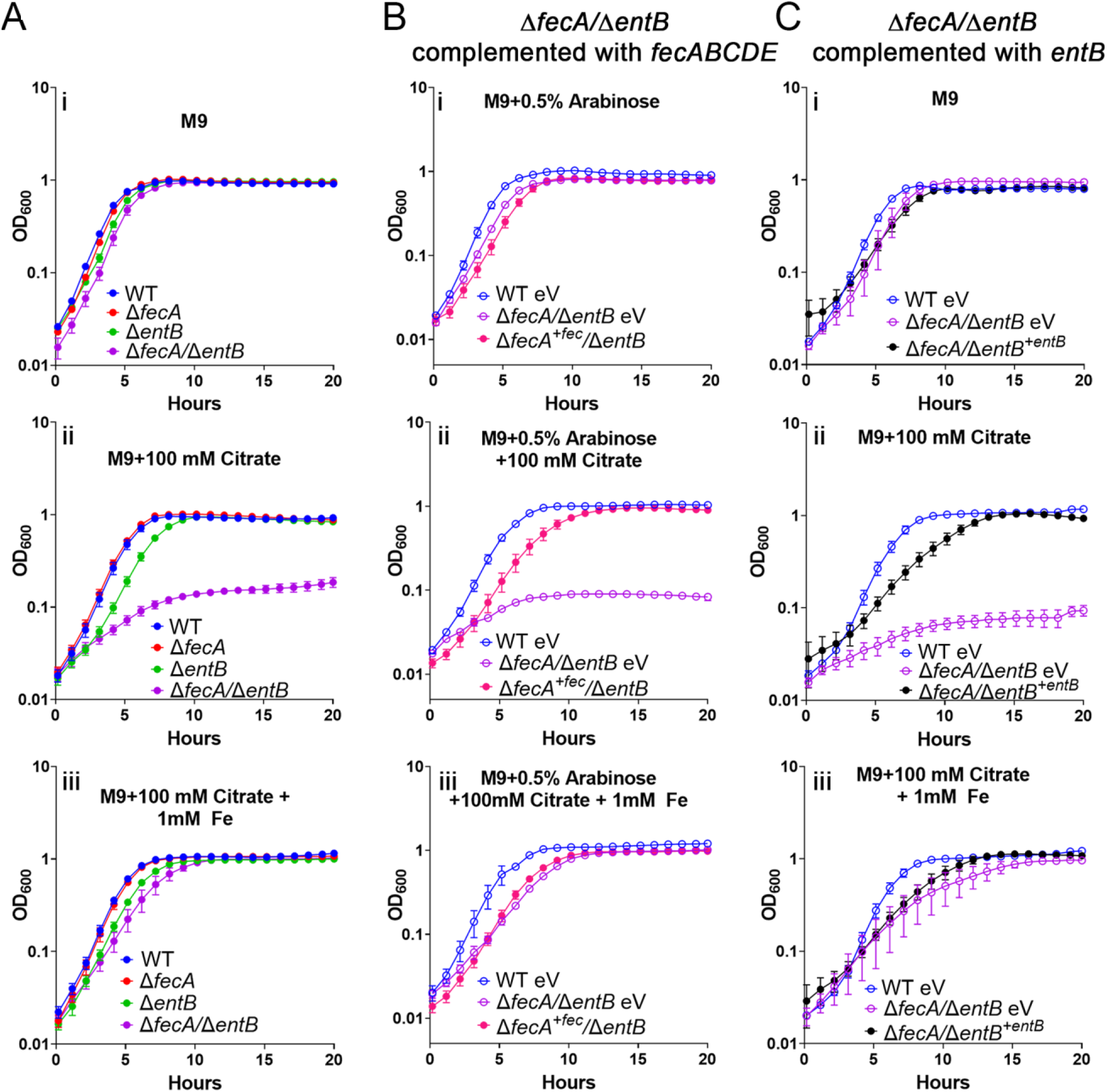
HM7 uses ferric citrate as an iron source through the *fec* system and enterobactin. Growth in M9 medium (i), M9 medium supplemented with 100 mM citrate (ii), and M9 medium supplemented with both 100 mM citrate and 1 mM FeCl_3_ (iii) of (A) WT HM7, single mutants Δ*fecA*, Δ*entB* and double mutant Δ*fecA*/Δ*entB*, (B) WT HM7 expressing empty pBAD vector (WT eV), and Δ*fecA/*Δ*entB* expressing empty pBAD vector (Δ*fecA/*Δ*entB* eV), and Δ*fecA*/Δ*entB* complemented with *fecABCDE (*Δ*fecA*^*+fec*^*/*Δ*entB*). All media in these conditions are supplemented with 0.5% arabinose to induce expression. (C) WT HM7 expressing empty pGEN vector (WT eV), and Δ*fecA*/Δ*entB* expressing empty pGEN vector (Δ*fecA/*Δ*entB* eV), and Δ*fecA*/Δ*entB* complemented with *entB* under control of its native promoter (Δ*fecA/*Δ*entB*^*+entB*^). 0.4% glucose was used as the sole carbon source in all conditions. Growth curves show averages of three to five biological replicates, error bars are SEM.

To establish that HM7 could specifically use the *fec* system to acquire iron via ferric citrate, we took a genetic approach, complementing the Δ*fecA*/Δ*entB* double mutant with each single system. Unsurprisingly, growth of the Δ*fecA*/Δ*entB* double mutant was rescued by genetic complementation with *entB* (**Fig. 3Ci, ii**, and **iii**). However, *fecABCDE* was also sufficient to rescue growth (**Fig. 3Bi, ii**, and **iii**). *fecA* was not sufficient to rescue growth, indicating that the Δ*fecA* mutant is a polar mutation, although that does not change the interpretation of our previous results.

### Ferric citrate uptake is an *in vitro* fitness factor when HM7 cannot utilize enterobactin

The association of *fecA* with UPEC strains (**Fig. 2B**) and HM7 using ferric citrate as an iron source (**Fig. 3**) indicates that the presence of the *fec* system could provide UPEC with a competitive advantage. Initially we assessed growth of WT HM7, and the three mutants, Δ*fecA*, Δ*entB*, Δ*fecA*/Δ*entB*, in pooled *ex vivo* urine (**Fig. S3**). Surprisingly there seemed to be no significant growth defect in any of these mutants compared to WT. Therefore, we turned to a more sensitive assay to assess the advantage the *fec* could provide and performed competition experiments in pooled human urine. We tested WT against the Δ*fecA* mutant and observed that there was no competitive disadvantage of the mutant strain compared to WT (**Fig. 4A**). Both strains could still use enterobactin, indicating that perhaps the siderophore is the preferred mechanism to acquire iron. This was confirmed when competing WT and Δ*entB*; the mutant had a subtle, but significant disadvantage (**Fig. 4A**). This disadvantage was exacerbated and trended towards significance when WT was competed against the Δ*fecA*/Δ*entB* double mutant (**Fig. 4A**), indicating that both systems contribute to the survival of HM7, but the function of enterobactin masks the role of *fec*.

**Figure 4:**
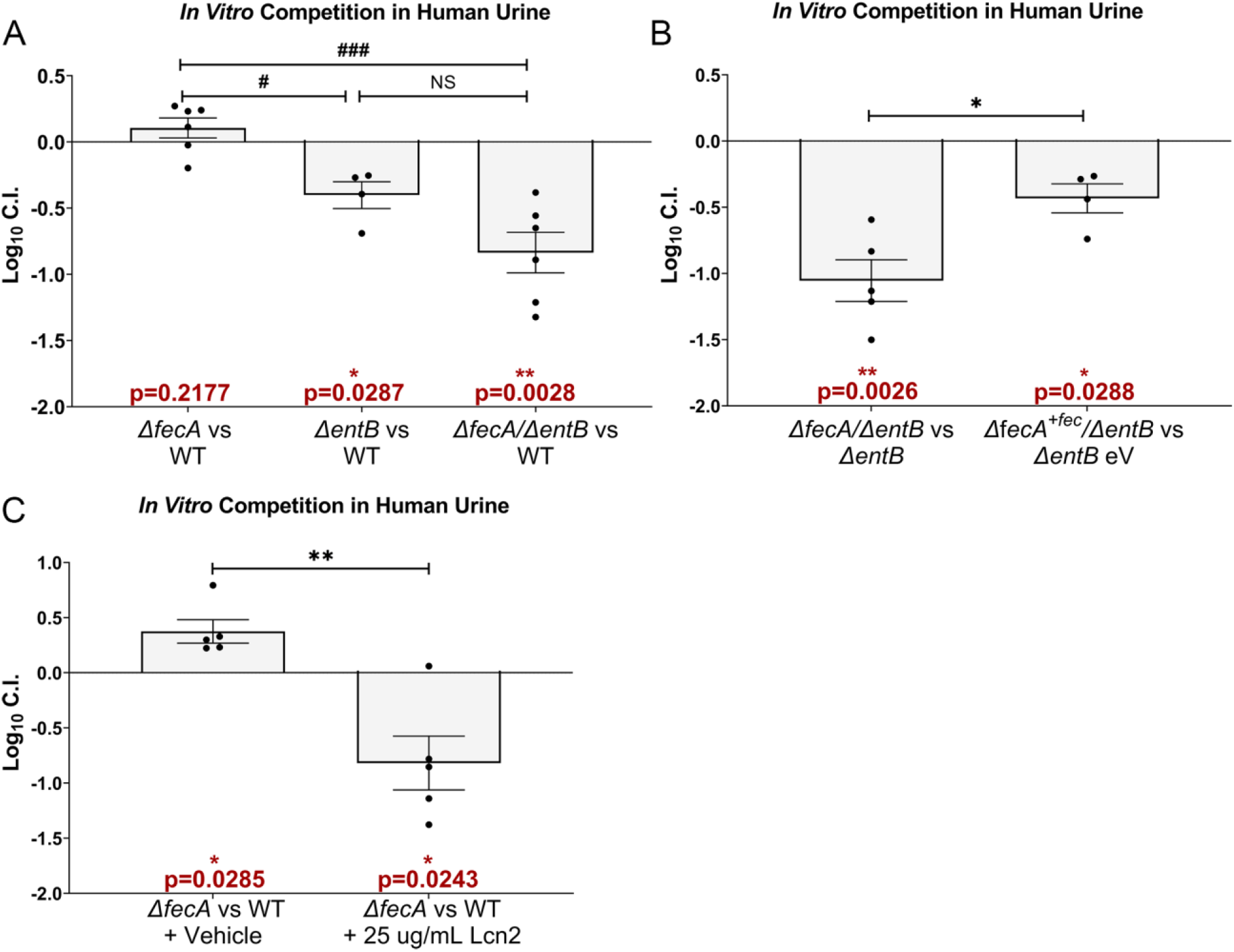
Ferric citrate uptake is an *in vitro* fitness factor in the absence of enterobactin. *In vitro* fitness of strains or conditions were determined in *ex vivo* pooled human urine. All strains were inoculated in a 1:1 ratio and grown for 24 hours at 37°C with aeration and their log_10_ competitive index (C.I.) determined. A log_10_ C.I. <0 indicates the first listed strain was outcompeted by the second. (A) Indicated strains were competed. (B) The Δ*entB* mutant expressing an empty vector (Δ*entB* eV) and the Δ*fecA/*Δ*entB* double mutant with the *fec* operon complemented in *trans* (Δ*fecA+*^*fec*^*/*Δ*entB*) were competed in urine supplemented with 0.5% arabinose and ampicillin (100 μg/mL). (C) WT HM7 was competed with Δ*fecA* and the urine was supplemented with either recombinant human lipocalin (Lcn2) or an equal volume of vehicle (25% glycerol). Red asterisks (*) indicate a significant competitive disadvantage, determined by one sample t-test, * p<0.05, ** p<0.005. Hashtags (#) compare log_10_ C.I. between indicated strains using ordinary one-way ANOVA with Sidak’s multiple test correction, # p<0.05, ### p<0.001. Black asterisks (*) compare log_10_ CI between indicated strains or conditions using unpaired t test, * p<0.05, ** p<0.005. Bars indicate mean, error bars are ±SEM, each dot represents an independent experiment.

To dissect the precise contribution of the *fec* system, we competed the Δ*entB* mutant with the Δ*fecA*/Δ*entB* double mutant. The double mutant had a significant defect (**Fig. 4B**). This defect is largely specific to the *fec* system since complementing the double mutant with *fecABCDE* was sufficient to partially rescue the defect (**Fig. 4B**). In the urinary tract during infection, Lcn2 counteracts enterobactin. Therefore, to mimic the host’s infectious environment, we added recombinant Lcn2 to these competitions. We determined that 25 μg/mL of Lcn2 was sufficient to inhibit HM7 growth in an iron-limited environment (**Fig. S4**) and then supplemented that amount to pooled human urine and competed WT and the Δ*fecA* mutant. With the addition of Lcn2, the Δ*fecA* mutant now had a significant competitive disadvantage (**Fig. 4C**). This provides further evidence that in the absence or inhibition of enterobactin the *fec* system is a fitness factor.

### Ferric citrate uptake is an *in vivo* fitness factor

Finally, we wanted to determine if the *fec* system was an *in vivo* fitness factor. Using the ascending UTI mouse model, we co-infected female CBA/J mice with WT and the Δ*fecA* mutant, allowed the infection to progress for 48 hours and harvested the urine, bladder, and kidneys to calculate log_10_C.I. The Δ*fecA* mutant had a significant disadvantage in all three organ sites (**Fig. 5**), definitively defining it as a fitness factor.

**Figure 5:**
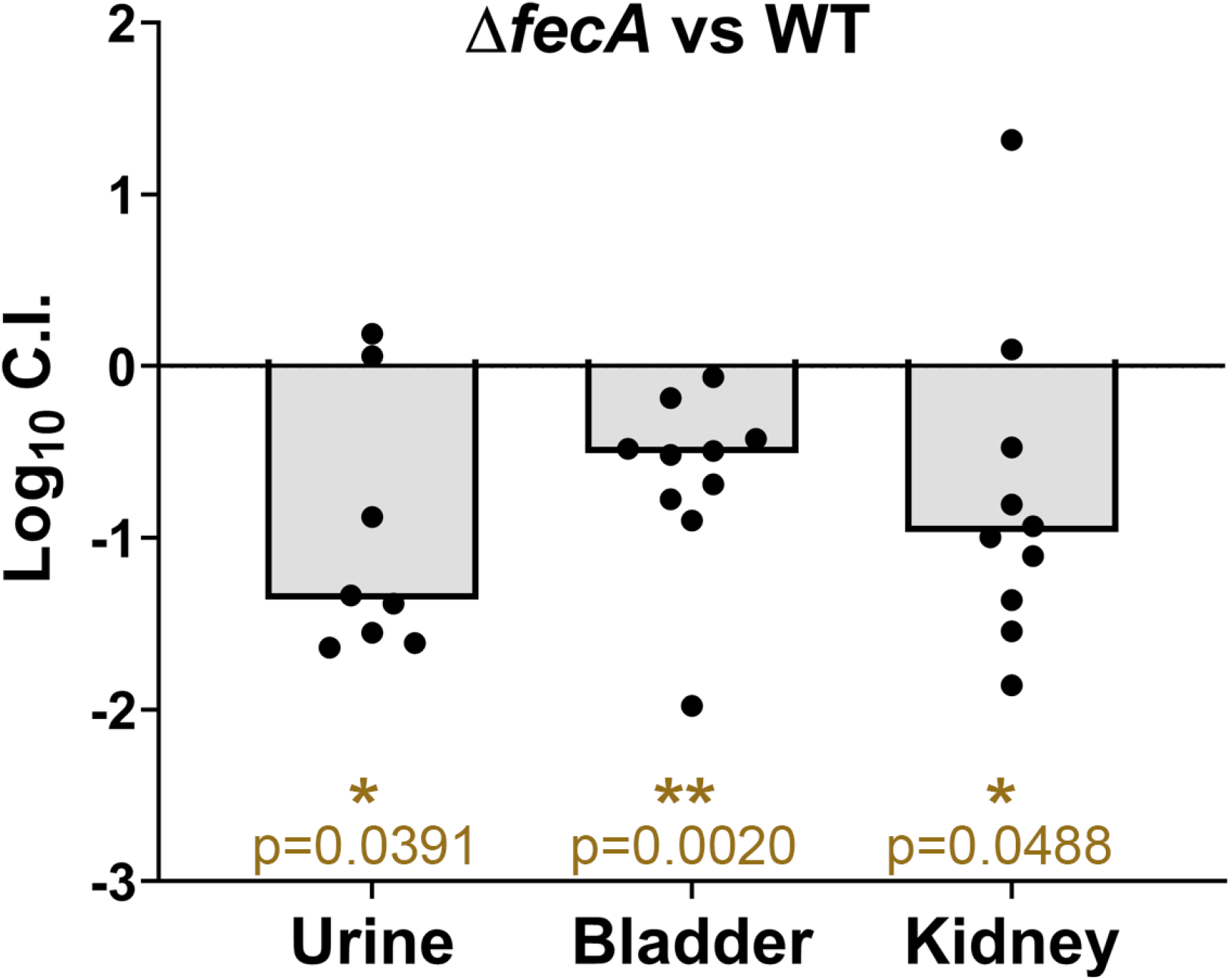
Ferric citrate uptake is an *in vivo* fitness factor. WT HM7 and single mutant Δ*fecA* were combined in a 1:1 ratio and transurethrally inoculated into CBA/J mice. Competitive indices were calculated 48 hours post infection. Symbols are individual animals, bars are median. Significance was determined with Wilcoxon signed-rank test, * p<0.05, ** p<0.005

We hypothesized that the Δ*fecA* mutant had a defect *in vivo*, due to the presence of Lcn2, as we saw in our *in vitro* competitions that were supplemented with Lcn2 (**Fig. 4C**). Lcn2 is highly elevated in the bladders and kidneys of mice infected with WT HM7 (**Fig. S5A, B**). Lcn2 levels correlated with increased CFU burden in the kidneys, where it is produced (37) (**Fig. S5C**). To determine if Lcn2 was responsible for the competitive disadvantage of the Δ*fecA* mutant, we repeated the competition experiments with Lcn2 knock-out mice (*Lcn2*^-/-^). However, the *Lcn2*^-/-^ mice are in a different genetic background, C57BL/6, rather than CBA/J, so we repeated the competition in the WT (C57BL/6) mouse background as well. While there was a subtle difference in the log_10_C.I. of the bladders between WT and Lcn2^-/-^ mice that was trending towards significance, the Δ*fecA* mutant no longer had a disadvantage compared to WT in the C57BL/6 background (**Fig. S6**). This discrepancy in results indicates the differences between the mouse strains. Overall, we conclude that ferric citrate uptake through the *fec* system is a *bona fide* fitness factor in UPEC strain HM7, allowing it to acquire iron from the host in a manner not inhibited by Lcn2.

## Discussion

Iron acquisition is an essential virulence factor in UPEC, because most iron in the host is sequestered. Subsequently, UPEC relies on specific iron acquisition systems such as siderophores or heme receptors to scavenge iron from otherwise inaccessible sources and these systems are essential for UPEC pathogenesis (10, 38, 39). Our work shows there is another understudied and overlooked iron acquisition system that enhances UPEC pathogenesis: ferric citrate uptake, encoded by the *fec* system.

Our study focuses on a recent clinical UPEC isolate, HM7. This strain encodes a sole siderophore, enterobactin, and lacks the functional redundancy in iron acquisition systems normally observed in UPEC strains. We assumed that HM7 was employing another method to acquire iron from the host and used RNA-seq to define its iron regulon. Under iron-limiting conditions, we found almost every component of ferric citrate uptake (*fecABCE* and *fecIR*) was highly and significantly upregulated (**Table 1, Table S2, Fig. 2A)**. Interestingly, *fecD* was not highly upregulated. While the rest of the genes in the system had log_2_ fold-change (FC) values ranging from 2.6-5.1, *fecD* had a log_2_FC of 0.8, and unlike the rest, this change was not significant. This is intriguing given that *fecD* is the second to the last gene in the operon, and yet the gene after it, *fecE*, is significantly and highly upregulated. *fecD* and *fecC* encode the permeases of the transport system that form a channel in the inner membrane of the bacterium (24, 40). Permeases can form homodimers, or heterodimers, and it is tempting to speculate the modest upregulation of *fecD* indicates that there is a preference for FecC homodimers as opposed to FecC/FecD heterodimers. Potentially, there could be an alternative start site that *fecE* utilizes, explaining its higher expression levels. *fecE* encodes the ATPase of this system (40), which is essential for activity of this ABC transporter. However, precisely defining this mechanism will require future studies.

We uncovered a strong association of the *fec* system with UPEC strains compared to fecal or environmental strains, another indication that the *fec* system is a virulence factor. (**Fig. 2B**). Given how common the *fec* system is within UPEC and MPEC, it could be a virulence factor in other pathogenic *E. coli*. For example, the citrate levels in plasma vary from 100-150 μM (41) and while these levels are lower than in urine or milk, they are still sufficient for robust upregulation of *fecA* (**Fig. 2D**). Potentially the *E. coli* that cause bloodstream infections could also utilize *fec* to acquire iron. In fact, a recent study exploring conjugative plasmids in pathogenic *E. coli* found a plasmid that encoded the *fec* system conferred a modest *in vivo* competitive advantage during bacteremia (42). This was also tested in the UTI model, and when this plasmid was conjugated into a different *E. coli* strain loss of *fec* resulted in an extremely mild reduction in fitness (log_10_C.I. ∼-0.1 in the bladder, and ∼-0.2 in the kidneys). However, this result could not be recapitulated in its parent strain. Other iron acquisition systems in these strains were not defined and could explain the divergence of results, demonstrating how functional redundancy of iron acquisition systems can mask the contributions of specific systems.

The *fec* system seems lower on the hierarchy of iron acquisition systems but becomes more important the fewer iron acquisition systems a strain produces. In a large cohort of UPEC strains, about ∼9% encoded a single traditional iron acquisition system, enterobactin, like HM7. While a relatively small percentage, it is still a part of the population that would likely rely more heavily on a system like *fec*, especially given that enterobactin is not highly effective in the urinary tract (38). This seems to be case since the *fec* system in these isolates is more enriched compared to more traditional UPEC strains (**Fig. 2C**). Interestingly, a very small population of these strains (2%) seemed to encode none of the traditional iron acquisition systems. However, the sequencing quality of these genomes is quite poor, and follow up studies are needed to confirm these results. If these results are confirmed, these strains could be an excellent tool to discover additional novel or understudied iron acquisition systems.

*In vitro* competition in pooled human urine showed that the *fec* system provides a competitive advantage, but this advantage is contingent on the absence of enterobactin (**Fig. 4**). Enterobactin seems to be the preferred method of iron acquisition *in vitro*, since loss of enterobactin is sufficient to cause a small reduction in fitness (**Fig. 4A**). Furthermore, addition of Lcn2 was sufficient to inhibit the function of enterobactin, allowing the *fec* system to provide an advantage (**Fig. 4C**). Lcn2 is present in high levels in the urinary tract during infection (43); therefore, these *in vitro* competitions with the addition of Lcn2 are likely a closer representation of UTI.

The gene expression profile of UPEC during CBA/J mouse infection closely mimics the UPEC transcriptome during human infection (44). Therefore, we are reasonably confident that the results from the mouse model are relevant to human infection. When WT HM7 was competed against the *ΔfecA* mutant, the mutant had a disadvantage in the urine, bladder, and kidneys (**Fig. 5**). While this result is different than the *in vitro* competition in human urine alone, it aligns with the *in vitro* competitions supplemented with Lcn2 and implies that the *fec* system provides an advantage *in vivo* because HM7 is unable to use enterobactin. This is likely caused by Lcn2; mice infected with HM7 had increased production of Lcn2 in the bladder and kidneys (**Fig. S5**).

We attempted to confirm this hypothesis using *Lcn2*^-/-^ mice. If Lcn2 is essential for the competitive advantage of the *fec* system that advantage should be abrogated in the knock-out line. The *Lcn2*^-/-^ mice were in a C57BL/6 background, therefore, we re-tested WT HM7 and *ΔfecA* mutant in WT C57BL/6 mice. Unfortunately, there was no loss in fitness in the *ΔfecA* mutant in WT C57BL/6 mice (**Fig. S6**). However, there are several genetic differences between these mouse lines (45) that could account for these differences. For example, of the 10 CBA/J mice we infected, 100% of them had kidney colonization, while only 35% of the 20 C57BL/6 mice had kidney colonization. While the contribution of Lcn2 to the mechanism of ferric citrate uptake via the *fec* system has not been definitively proven, it seems a promising explanation.

In summary, we have uncovered a novel mechanism by which UPEC acquires iron from the host via ferric citrate uptake. During UTI Lcn2 is highly produced, blocking the usage of enterobactin. In response, UPEC uses the *fec* system to import ferric citrate present in the urinary tract as an iron source (**Fig. 6**). The *fec* system is highly prevalent in UPEC strains and is yet another instrument in its highly diverse arsenal to survive within the harsh environment of the urinary tract.

**Figure 6:**
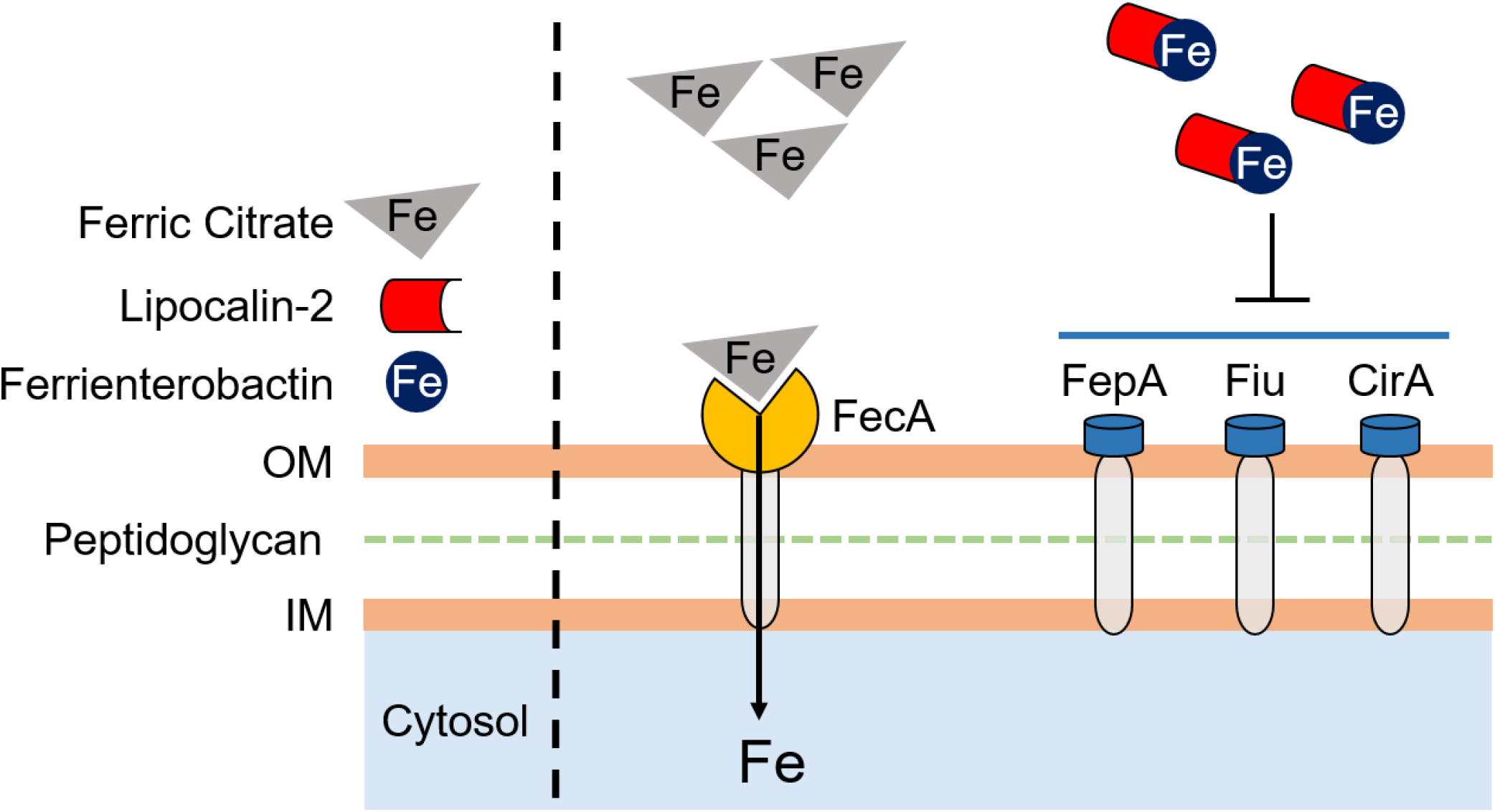
Model of UPEC utilization of ferric citrate.

## Acknowledgments

This work was supported by Public Health Service grant R01AI059722 from the National Institutes of Health. We would also like to thank members of Dr. Michael Bachman’s lab, specifically, Dr. Jay Vornhagen for providing us with the expression plasmid for Lcn2 purification and Dr. Caitlyn L. Holmes for both the WT and *Lcn2*^-/-^ C57BL/6 mice.

## Methods

### Bacterial culture conditions, growth curves, mutant construction, and complementation

Clinical UPEC isolate HM7 was routinely cultured at 37°C with aeration in LB, M9 medium supplemented with glucose, or filter-sterilized pooled human urine. Mutant and complemented strains were cultured with antibiotics. Mutants were constructed using lambda red mutagenesis and complementation vectors constructed with Gibson assembly. For a detailed description, refer to **Text S1**.

### Chrome Azurol S Assay

Chrome Azurol S (CAS)-agar was prepared as defined in (31). Strains were cultured overnight with aeration at 37°C in LB with appropriate antibiotics. Five μL of the overnight culture was spotted onto the CAS-agar plate, and then incubated overnight at 37°C. The next morning the plates were imaged using Qcount Software.

### RNA isolation and library preparation, and sequencing

*E. coli* HM7 was cultured overnight in M9 medium, supplemented with 0.4% glucose, shaking at 37°C. Overnight cultures were diluted 1:100 in M9 medium with 0.4% glucose supplemented with either 36 μM FeCl_3_ (Sigma), or 150 μM 2,2’ Dipyridyl (Sigma) and grown to mid-log phase (0.4-0.6 OD_600_). Cultures were then treated with Bacterial RNA Protect (Qiagen), harvested by centrifugation and the pellets stored at -80°C. This was performed in biological triplicate. RNA was isolated using a similar method from (28, 44). The libraries were prepared using NEBNext Ultra II Directional RNA Library Prep Kit and sequenced using an Illumina NextSeq-500 (paired end, 38 bp read length). For a detailed description, refer to **Text S1**.

### Genome assembly, RNA-seq data processing and differential expression analysis

Raw sequencing data was preprocessed using BBTools (38.18) (46). BBDuk was used to remove Illumina adapter sequences, and to quality trim and filter the reads (minlength=20, trimq=14, maq=20, maxns=1). The HM7 genome was re-assembled based on sequencing from (28) using Flye long read assembler (47) with Trestle repeat resolve parameter on then the quality controlled reads were aligned to the HM7 genome using BWA (0.7)(48). The resulting alignment files were filtered (mapping quality > 10) using samtools (1.11) (49) with counts for each feature were generated using htseq-count (0.13.5) (50). Alignment details shown in **Table S3**. Differential expression analysis was performed using R package DESeq2 (51).

### qRT-PCR

Strains were grown to mid-log cultures and RNA isolated as described above and reverse-transcribed into cDNA using iScript (Biorad). qRT-PCR was performed on a QuantStudio 3 PCR machine (Applied Biosystems) using PowerUp Syber Green mastermix (Applied Biosystems). For a detailed description, refer to **Text S1**.

### Purification of Lipocalin-2

Recombinant human Lipocalin-2 (Lcn2) expressed as a glutathione S-transferase (GST) fusion protein (52) (a kind gift from Dr. Michael Bachman) in XL-1 Gold *E. coli* protein was purified in a similar manner as previously described (53, 54). For a detailed description, refer to **Text S1**.

### *In vitro* growth competition

Strains were cultured overnight in M9 medium supplemented with 0.4% glucose at 37°C with aeration and appropriate antibiotic selection. The next day, the OD_600_ was determined for each strain, the strains were OD_600_ -matched, and then diluted 1:100 into 3 mL of pooled human urine. Where applicable, Lcn2 was added to a final concentration of 25 μg/mL, or the vehicle control (25% glycerol) in an equal volume. 0.5% arabinose (final concentration) induced the complemented strains, and ampicillin added to maintain the plasmid. Input CFUs were determined for each strain through drip plating of serial dilutions on plain LB agar and antibiotic selection (chloramphenicol or kanamycin). The strains were then grown overnight, shaking at 37°C, and the output CFU of each strain determined in the same manner as the input.

C.I. is a ratio of the input versus the output and is calculated as follows:

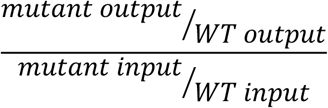

Log_10_CI <0 indicates that the WT outcompetes the mutant, and a log_10_CI >0 indicates the mutant outcompetes the WT. When competing Δ*entB* and Δ*fecA*/Δ*entB*, Δ*entB* was “WT”, and Δ*fecA*/Δ*entB* was “mutant”. When competing Δ*entB* eV and Δ*fecA*^*+fec*^/Δ*entB*, Δ*entB* eV was “WT”, and Δ*fecA*^*+fec*^/Δ*entB* was “mutant”.

### Murine UTI model

We used three different mouse strains: CBA/J, C57BL/6 WT and C57BL/6 *Lcn2*^-/-^ (55). CBA/J mice were purchased from Jackson Laboratories, while both the C57BL/6 WT and C57BL/6 *Lcn2*^-/-^mice were a kind gift from Dr. Michael Bachman and bred in-house. All mice used were female. For a detailed description, refer to **Text S1**.

## Data accessibility

Data available on in NCBI’s Gene Expression Omnibus repository under accession number GSE188170.

## Supplemental Figures

**Supplemental Figure 1.**
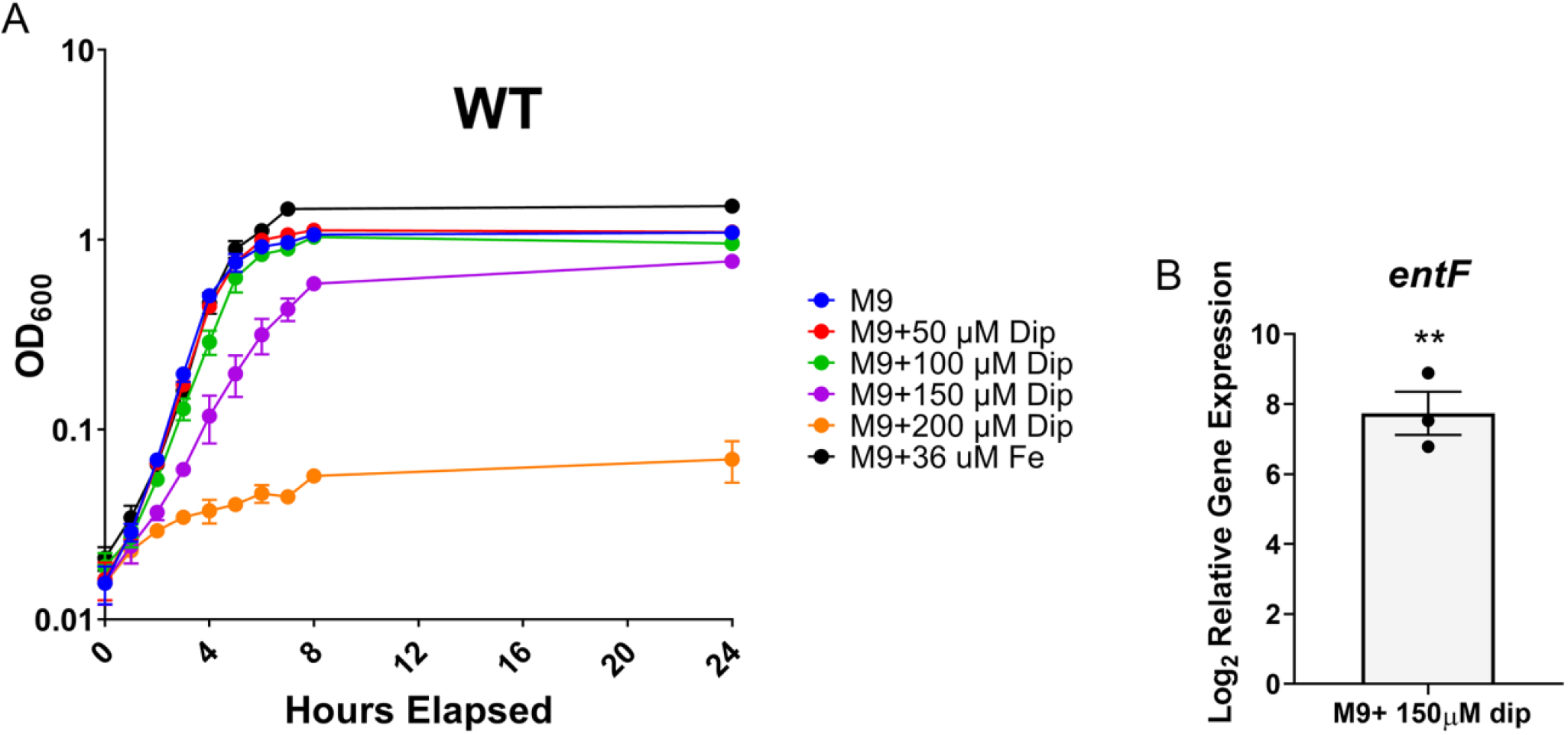
(A) Growth of WT HM7 in M9 medium with 0.4% glucose as the sole carbon source (M9), as well as supplemented with 36 μM FeCl_3_ or increasing amounts of the iron chelator 2,2’-dipyridyl (Dip). WT HM7 was cultured overnight in M9, and then subcultured 1:100 into 3 mL medium in culture tubes and grown at 37°C with aeration. OD_600_ was measured on a plate reader for eight hours, taking a reading every hour, and then another reading was taken at 24 hours. Results are an average of three to four biological replicates, error bars represent ±SEM. (B) Gene expression of *entF* in WT HM7 in M9 medium supplemented with 150 μM Dip compared to M9 supplemented with 36 μM FeCl_3_. Gene expression was assayed through qRT-PCR. Bars are the average of three biological replicates, dots are the values from each biological replicate and error bars are ±SEM, and asterisks indicate significant upregulation, determined by one sample t-test, ** p<0.01.

**Supplemental Figure 2.**
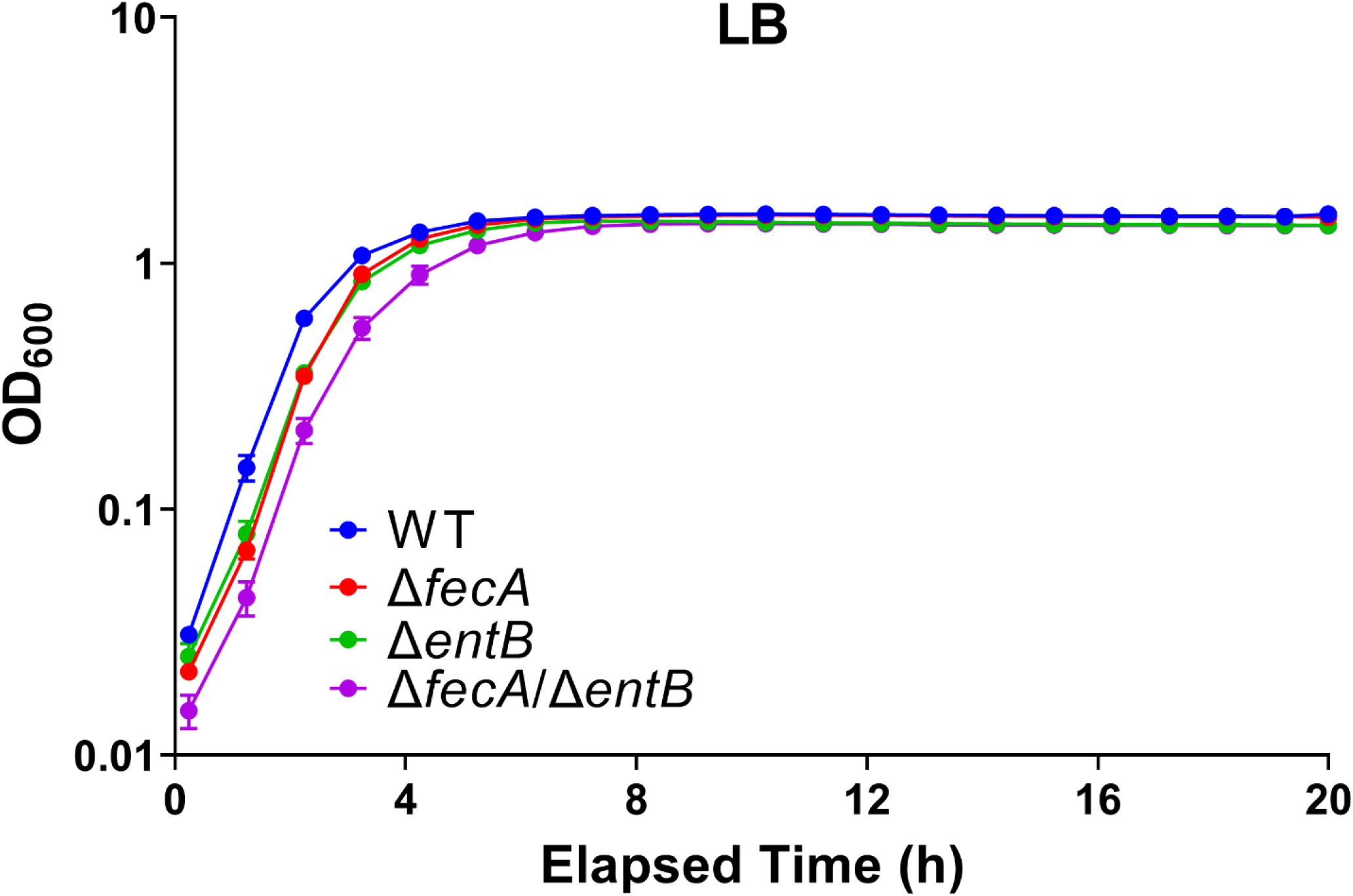
Growth of WT HM7, Δ*fecA*, Δ*entB*, and Δ*fecA/*Δ*entB*, in LB. Results are an average of four to five biological replicates, bars represent ±SEM.

**Supplemental Figure 3.**
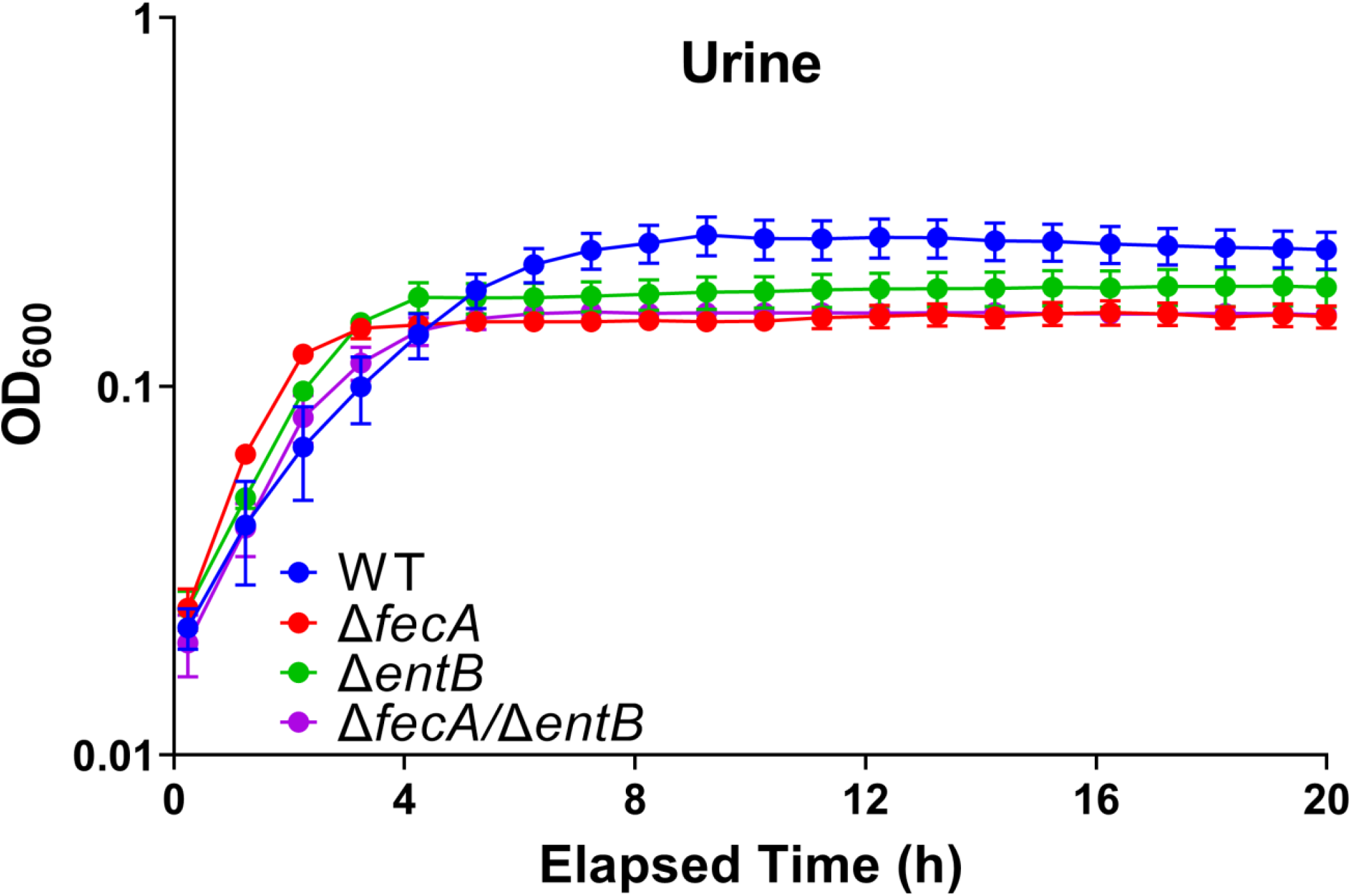
Growth of WT HM7, Δ*fecA*, Δ*entB*, and Δ*fecA/*Δ*entB* in *ex vivo* urine pooled from healthy female volunteers. Results are an average of four to five biological replicates, bars represent ±SEM.

**Supplemental Figure 4.**
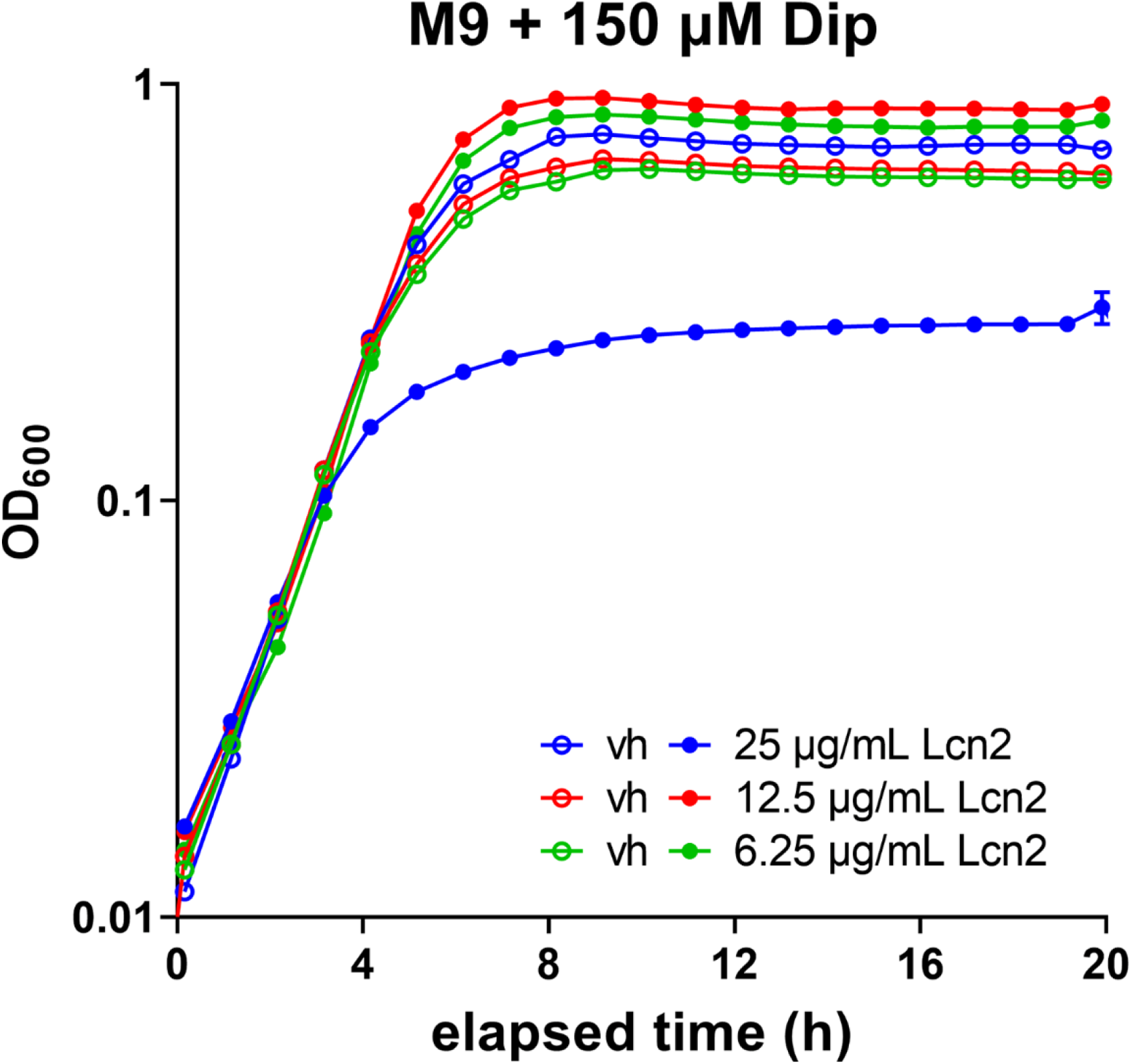
Growth of WT HM7 supplemented with recombinant human lipocalin (Lcn2). WT HM7 was grown in an iron starved state (M9 medium supplemented with 150 μM Dip) with increasing amounts of Lcn2. An equal volume of the vehicle (vh, 25% glycerol) for each amount was added as a control. Results are an average of two biological replicates, bars represent ±SEM.

**Supplemental Figure 5.**
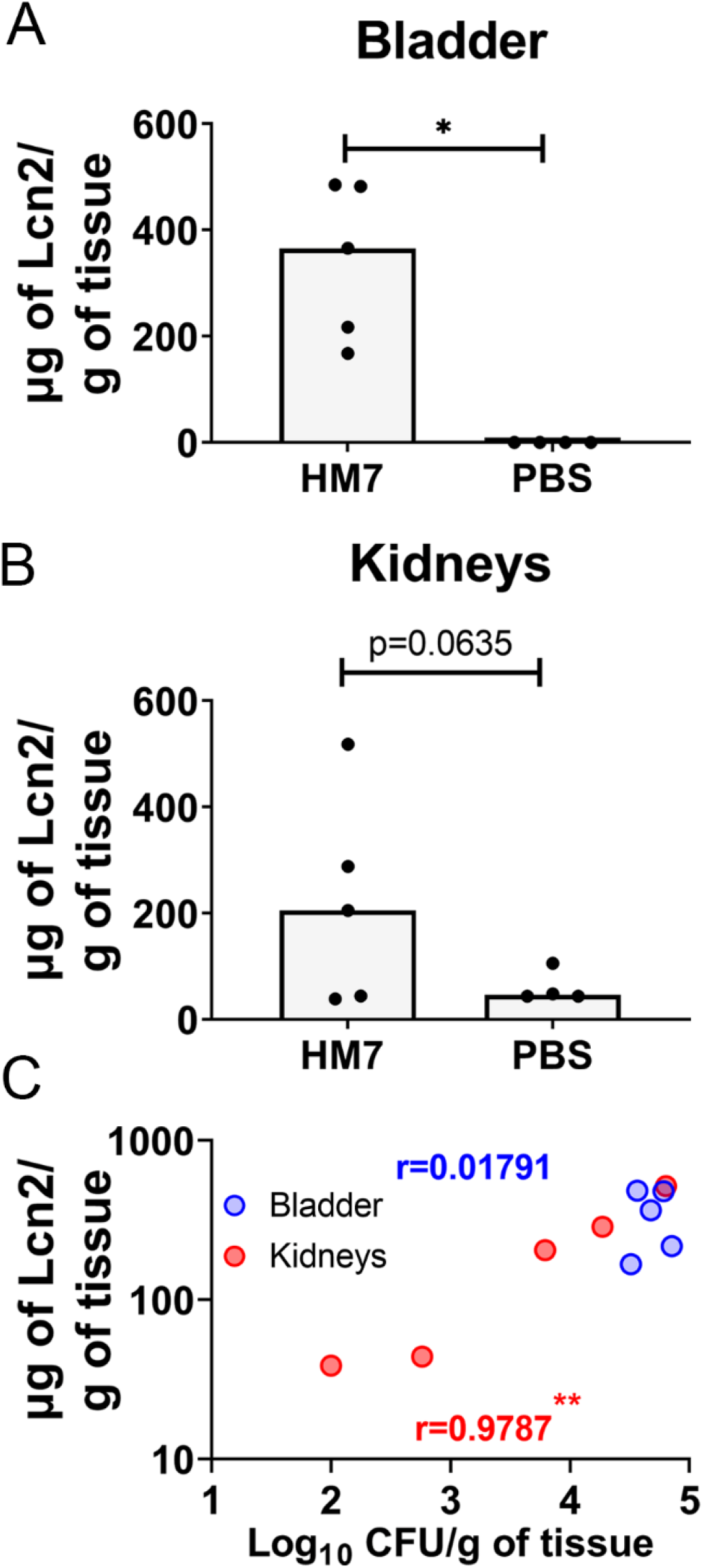
Quantification of lipocalin (Lcn2) production during murine infection. CBA/J mice were infected either with WT HM7 or mock infected with PBS. Lcn2 levels were quantified via ELISA in the (A) bladder and (B) kidneys. (C) Lcn2 levels were plotted against CFU burden of mice infected with HM7. Pearson correlation coefficient (r) for bladder is displayed in blue, and for kidneys is displayed in red. Dots indicate individual mice, bars are median. Significance was determined via Mann-Whitney test, * p<0.05.

**Supplemental Figure 6.**
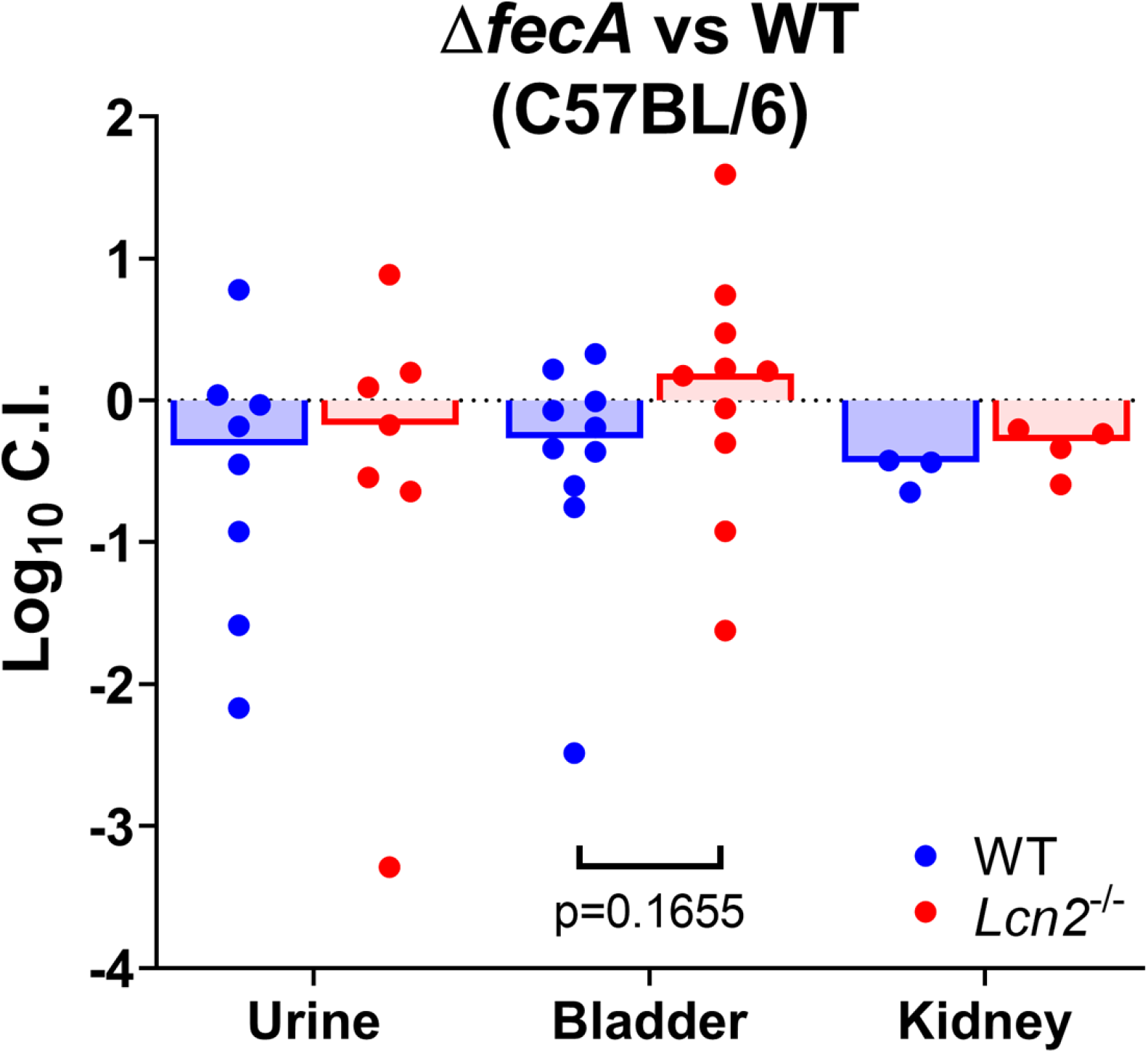
WT HM7 competed with Δ*fecA* in WT C57BL/6 mice (WT) and lipocalin null (*Lcn2*^-/-^) mice. Competitive indices were calculated 48 hours post infection. Historically, C57BL/6 mice have poor kidney colonization; only 3/10 WT mice and 4/10 *Lcn2*^-/-^ mice had detectable CFU in the kidney. Dots are individual animals, bars are median. Log_10_CIs were compared using Mann-Whitney test.

